# DIELECTRIC CONTINUUM MODELING FOR THE DE NOVO DESIGNED CALCIUM CHANNEL CALC6_3: STRUCTURE PREPARATION, MESH GENERATION, AND CALCIUM SELECTIVITY

**DOI:** 10.64898/2026.10.20.760449

**Authors:** Dexuan Xie, Robert S. Eisenberg

**Affiliations:** Department of Mathematical Sciences, University of Wisconsin-Milwaukee, Milwaukee, WI 53201, USA; Department of Applied Mathematics, Illinois Institute of Technology, Chicago, IL, USA; Department of Physiology and Biophysics, Rush University Medical Center, Chicago, IL, USA

**Keywords:** CalC6_3 calcium channel, pore-preserving Gaussian molecular surface mesh, size modified Poisson–Boltzmann finite element solver, calcium selectivity, integrated ion-channel simulation package

## Abstract

This paper presents an integrated software package for continuum dielectric studies of the de novo designed calcium channel CalC6_3. The package combines Python scripts for cleaning and repairing the original PDB structure (PDB ID: 9DZW) with PDB2PQR to generate a corresponding PQR file containing atomic coordinates, partial charges, and radii. It incorporates the Gaussian Ion Channel Molecular Surface Generation package to construct high-quality triangular molecular surface meshes that preserve the complex channel pore geometry while maintaining sufficiently smooth exterior molecular surfaces. These surface meshes are used to generate interface-fitted tetrahedral meshes of a simulation box consisting of protein, membrane, and solvent regions. The integrated software package further integrates the structure-preparation and mesh-generation procedures with the nonuniform size-modified Poisson–Boltzmann ion channel finite element solver to compute electrostatic potentials and ionic concentrations, streamlining the complete workflow from structure preparation to numerical solution. Numerical experiments for an ionic solution containing Ca^2+^, Na^+^, and Cl^−^ ions show preferential Ca^2+^ accumulation in the narrow selectivity filter formed by six Glu residues, supporting the experimentally observed calcium selectivity of CalC6_3. They also suggest that a smoother molecular surface mesh can improve the convergence and performance of the finite element solver.

**AMS subject classifications:** 92-08, 65N50, 65N30

## 1. Introduction

Calcium ions play essential roles in a wide range of biological processes, including muscle contraction, neurotransmitter and hormone release, gene-expression regulation, and intracellular signal transduction. The controlled movement of Ca^2+^ across cell membranes is mediated by calcium-selective ion channels, whose selectivity filters must preferentially conduct Ca^2+^ in the presence of other physiologically abundant ions, particularly Na^+^ [4, 26]. Understanding the structural and electrostatic mechanisms underlying this remarkable selectivity is therefore an important objective in structural biology, biophysics, and computational biology.

Substantial progress has been made in determining high-resolution structures of native voltage-gated calcium (CaV) channels using cryogenic electron microscopy (cryo-EM). These structures have provided important insights into channel architecture, ion selectivity, gating, and pharmacological modulation. However, no CaV structure in an open state has yet been determined experimentally, possibly because the open state is labile, short-lived, and voltage-dependent. During protein purification and cryo-EM sample preparation, a controlled transmembrane voltage is generally absent, and activated channels are likely to enter closed or inactivated conformations [25]. For example, the cryo-EM structure of human CaV1.3 deposited under PDB ID 7UHG contains a closed intracellular gate and therefore cannot be used directly to simulate ion permeation [24]. This limitation has motivated computational efforts to construct putative open-state CaV models from experimentally determined closed-state structures.

Recently, Xue et al. constructed an open-pore model of CaV1.3 using a closed-state cryo-EM structure of CaV1.3 (PDB ID: 7UHG) and an open-state structure of the eukaryotic sodium channel Na_V_Eh (PDB ID: 7X5V) as a template for the open-pore model [23, 27]. Specifically, the four S6 helices of Na_V_Eh were aligned with the corresponding S6 helices of CaV1.3, and template-derived position restraints were applied during molecular dynamics simulations to produce an open pore. Simulations using the resulting model showed a preference for Ca^2+^ over Na^+^ consistent with the expected valence selectivity of CaV1 channels [23]. This template-based construction is physically motivated by the evolutionary and structural relationships between voltage-gated sodium and calcium channels and provides a useful means of investigating CaV permeation when an experimentally determined open-state structure is unavailable. Nevertheless, the resulting pore is computationally constructed and has not been experimentally validated as an open-state pore of CaV1.3.

As an alternative to constructing an open-state model from a closed native calcium-channel structure, Liu et al. recently introduced a bottom-up strategy for designing calcium-selective channels with open pores [12]. Patch-clamp measurements showed that their tetrameric and hexameric channels had higher conductance for Ca^2+^ than for Na^+^, experimentally demonstrating preferential calcium conduction. Of particular interest is CalC6_3, a homohexameric channel formed by six symmetrically arranged subunits, each contributing a pore-lining helix to the central pore. Its narrow selectivity filter is formed by six Glu residues, confirming the motif of calcium selectivity first proposed by Nonner and Eisenberg [13]. The constriction has an estimated radius of approximately 3 Å, and the distance between diagonally opposed oxygen atoms in the filter is approximately 6.4 Å [12]. The CalC6_3 structure was determined by single-particle cryo-EM at a resolution of 3.75 Å and deposited in the Protein Data Bank under PDB ID 9DZW. The resolved structure is nearly identical to the computationally designed model, and provides an experimentally determined calcium-selective pore for structure-based simulations. Its reported pore geometry and dimensions also provide valuable reference data for assessing whether the molecular surface mesh required by dielectric continuum models preserves the geometric shape and dimensions of the channel pore. CalC6_3 therefore provides a useful model system for computational studies of calcium selectivity.

The CalC6_3 work is significant not only for de novo ion-channel design but also for computational studies of calcium selectivity. Unlike native CaV channels, CalC6_3 does not contain voltage-sensing domains or an activation gate that reproduces the complete gating mechanism of a native channel. It should therefore not be regarded as a structural substitute for the full CaV architecture. Instead, its open pore, experimentally demonstrated calcium preference, resolved cryo-EM structure, and relatively simple hexameric organization make it a valuable complementary channel model for investigating calcium selectivity using dielectric continuum models. In particular, CalC6_3 enables the study of how selectivity-filter geometry, protein atomic charges, and ionic sizes affect the distributions of Ca^2+^ and competing ions within the selectivity filter. Many coarse-grained versions of molecular dynamics are likely to depend on dielectric continuum models like those developed here. Such coarse graining seems unavoidable in molecular dynamics simulations involving physiological intracellular Ca^2+^ concentrations as low as 10^−8^ mole per liter (mol/L).

In dielectric continuum models, the channel protein, membrane, and surrounding solvent are represented as dielectric media, while mobile ions are described by continuous concentration functions. Among these models, the Poisson–Boltzmann ion channel (PBic) model and its size-modified variants describe the electrostatic potential and ionic concentrations at thermodynamic equilibrium, where the net flux of each ionic species is zero [9, 17, 19]. In contrast, the Poisson–Nernst–Planck ion channel (PNPic) model and its size-modified variants describe the electrostatic potential, ionic concentrations, and ionic fluxes under nonequilibrium electrodiffusion [20, 21, 22]. Compared with molecular dynamics simulations [3, 14], PBic and PNPic models can substantially reduce computational cost and efficiently treat complex ionic solutions. They may also give insight into general principles of calcium and sodium selectivity used by evolution in many proteins and contexts. They are therefore useful for investigating electrostatic mechanisms, ionic distributions, ion transport, and structure–function relationships in ion channels.

To study CalC6_3 using dielectric continuum models, the original 9DZW PDB file must first be processed because it contains 6,550 hydrogen atoms and a `CRYST1` record inherited from the in silico design and structure-determination procedure, while lacking 175 heavy atoms. We removed the deposited hydrogen atoms and the `CRYST1` record and transformed the molecular structure into a Cartesian coordinate system whose origin is located at the center of the channel and whose *z*-axis is aligned with the membrane-normal direction. This coordinate system is required for constructing the protein-membrane-solvent simulation box used by our dielectric continuum ion channel models. Something like this model is needed in most coarse grained treatments of molecular dynamics, particularly those that restrict the motions of some of the atoms of a channel protein. We subsequently converted the processed PDB file into a PQR file using the PDB2PQR software package [6]. During this conversion, PDB2PQR repaired the missing heavy atoms, determined the protonation states at pH 7.0, added the corresponding hydrogen atoms, optimized their orientations for hydrogen bonding, and assigned the atomic partial charges and radii according to the AMBER force field. The PQR file provides the atomic data required for constructing the triangular molecular surface mesh, a key step in generating simulation-box finite element meshes.

Generating a triangular molecular surface mesh from the PQR file is challenging because CalC6_3 contains a channel pore with a complex geometry and a narrow calcium-selectivity filter. The quality of this surface mesh is critical because it defines the boundary of the protein region and the dielectric interfaces separating the protein region from the membrane and solvent regions. Artificially displacing atoms near the pore to facilitate molecular surface generation may substantially distort the protein environment and electric field. It distorts the experimentally determined channel-pore structure and compromises its calcium-selectivity properties. Therefore, the pore geometry should be preserved without modifying atomic positions. At the same time, the molecular surface outside the pore region must remain sufficiently smooth for the generation of a high-quality finite element mesh.

Several molecular-surface definitions and computational packages are currently available for generating protein surface meshes. These include the solvent-accessible surface (SAS) [10], the solvent-excluded surface (SES) [15], the molecular-surface mesh generated by MSMS [16], and Gaussian molecular surfaces generated by TMSmesh [5, 7, 11]. However, in our tests, none of these methods produced a high-quality triangular surface mesh for CalC6_3 that both preserved the channel-pore geometry and satisfied the requirements of finite element calculations.

To address this difficulty, we recently developed the Gaussian Ion Channel Molecular Surface Generation (GICMSG) package for ion-selective channels with narrow selectivity filters [18]. In contrast to TMSmesh, which uses a single decay parameter throughout the molecular domain, GICMSG employs two spatially varying parameter functions to control exterior surface smoothness and channel pore geometry separately. This allows GICMSG to generate high-quality triangular meshes with smooth exterior surfaces while preserving essential pore geometry. However, TMSmesh fails to do so in our tests. In this work, GICMSG provides the molecular surface mesh of CalC6_3 used for subsequent volume mesh generation.

To generate a high-quality, interface-fitted finite element tetrahedral mesh for the simulation box, we integrated GICMSG into the mesh-generation package reported in [9]. The resulting package takes a PQR file as input and generates both the triangular molecular surface mesh and the interface-fitted tetrahedral mesh of the simulation box. This integration connects pore-preserving molecular surface generation with the construction of volume meshes required for dielectric continuum simulations of CalC6_3.

We further wrote Python scripts to clean and repair the original PDB file, `9DZW.pdb`, and to generate the required PQR file using PDB2PQR [6]. We then combined these scripts with the mesh-generation package and the nonuniform size-modified Poisson– Boltzmann ion channel (nuSMPBic) finite element package reported in [17], creating an integrated package for computing the electrostatic potentials, ionic concentrations, and electrostatic solvation free energies of CalC6_3. The integrated package takes the CalC6_3 PDB identifier as input and automates structure preparation, molecular surface and volume mesh generation, finite element solution of the nuSMPBic model, and visualization of the computed potential and concentration functions through color maps and two-dimensional plots. This integration reduces manual intervention and streamlines the complete workflow for CalC6_3 simulations.

Using the integrated package, we performed numerical experiments for CalC6_3 in an ionic solution containing 0.1 M NaCl and 0.1 M CaCl_2_. We generated two Gaussian molecular surfaces using GICMSG, one with a much smoother exterior surface than the other, while both closely preserve the channel pore geometry. The calculations produced the concentration functions of Ca^2+^, Na^+^, and Cl^−^, together with the electrostatic potential generated by the protein atomic charges and mobile ionic charges. These functions were used to characterize ionic distribution profiles across the membrane and examine preferential Ca^2+^ accumulation within the channel. The numerical results support the experimentally observed calcium selectivity of CalC6_3. They also suggest that a smoother molecular surface mesh can improve the convergence and performance of the nuSMPBic finite element solver. The complete calculation of one electrostatic potential function and three ionic concentration functions for CalC6_3 with 13,320 atoms, which involved solving a nonlinear finite element system with 141,915 unknowns, took approximately 44 seconds on a single CPU core of a Mac Studio equipped with an Apple M4 Max processor and 128 GB of memory, demonstrating the efficiency of the nuSMPBic finite element solver and the high performance of the integrated package.

Finally, we point out that previous Monte Carlo studies have investigated calcium selectivity using simplified models of L-type calcium channels, with idealized pore geometries and representations of glutamate-associated charged groups [1, 2]. In the present work, we use the nuSMPBic model to study CalC6_3 based on its experimentally determined structure [12], supplemented by reconstruction of missing atoms. Unlike the simplified models (see [13, Figure 1; Eq. (1) to (3)] for the one-dimensional Poisson-Nernst-Planck equations [1, 2] for the Monte Carlo simulation method), our computational domain is a simulation box consisting of protein, membrane, and solvent regions, with explicit molecular geometry and atomic charge distributions. The integrated package generates pore-preserving molecular surface meshes and interface-fitted tetrahedral meshes, enabling finite element calculations of electrostatic potentials and ionic concentrations while accounting for nonuniform ion sizes. This approach allows us to investigate calcium selectivity in the structural and dielectric environment of CalC6_3

The remainder of this paper is organized as follows. Section 2 presents the preparation of the structural data. Section 3 describes the generation of the triangular molecular surface mesh for CalC6_3. Section 4 presents the construction of the interface-fitted tetrahedral mesh of the simulation box. Section 5 reports the numerical test results and discussion, and Section 6 presents the conclusions.

## 2. Test data preparation

The atomic structure of the CalC6_3 calcium channel (PDB ID: 9DZW) was downloaded from the RCSB Protein Data Bank (https://www.rcsb.org). The original `9DZW.pdb`file contains 6,550 hydrogen atoms and one `CRYST1`record. It also lacks 175 heavy atoms from residue side chains distributed among the six subunits. The 9DZW structure was determined by cryogenic electron microscopy at a resolution of 3.75 Å and refined using an in silico design model [12]. The hydrogen atoms in the deposited structure were computationally generated rather than experimentally resolved. All deposited hydrogen atoms were therefore removed so that PDB2PQR [6] could add them consistently, assign protonation states at the specified pH 7.0, optimize hydrogen-bonding interactions, and assign atomic charges and radii. The `CRYST1`record was also removed because crystallographic unit-cell information was not required by the nuSMPBic finite element solver [17].

As required by the solver, the cleaned structure was translated and rotated so that the center of the channel pore was located at the origin and the pore axis coincided with the *z*-axis. The positive *z*-direction was defined as pointing from the intracellular side toward the extracellular side. The transformed structure was saved as `9DZW_modified.pdb`. Using PDB2PQR, we then generated `9DZW.pqr`from `9DZW modified.pdb`. The repaired CalC6_3 structure in `9DZW.pqr`provides a basis for computational investigations of calcium selectivity.

During PQR generation, PDB2PQR added the required hydrogen atoms, assigned atomic charges and radii at pH 7.0, and removed the water molecules stored in the PDB file. It detected 175 missing heavy atoms and reconstructed them using standard residue templates, local covalent-bonding information, and idealized side-chain geometries. It also reported `Unable to debump` warnings for five residues: ALA A57, ALA B57, LEU B71, ALA D57, and ALA F54. These warnings indicate that local structural optimization could not completely eliminate close atomic contacts involving these residues. The affected residues were retained, and the unresolved contacts were documented as limitations of the prepared structure. The computationally reconstructed coordinates should not be interpreted as experimentally resolved atomic positions. The resulting structure was visually inspected using VMD [8] and is shown in three views in Figure 2.1.

**Fig. 2.1:**
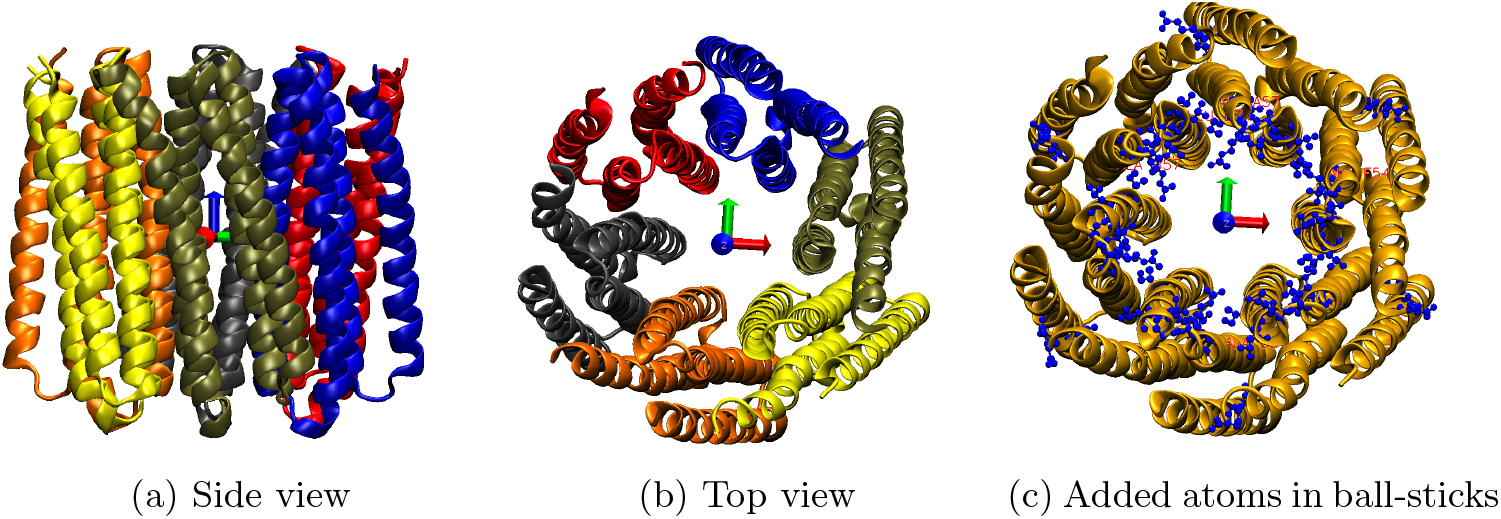
(a, b) Side and top views, respectively, of the repaired calcium channel structure represented by `9DZW.pqr`. The six identical subunits are shown in different colors and arranged symmetrically around the central pore. (c) Top view of the repaired structure, with the heavy atoms reconstructed by PDB2PQR shown as blue ball-and-stick representations and the six subunits shown in a gold cartoon representation.

Figures 2.1(a) and (b) show side and top views, respectively, with the six identical subunits displayed in different colors. The top view illustrates their approximately sixfold-symmetric arrangement around the central pore.

Figure 2.1(c) shows the heavy atoms reconstructed by PDB2PQR in blue using a ball-and-stick representation, with the six subunits shown in gold using a cartoon representation. Visual inspection indicates that most of the reconstructed atoms belong to residue side chains located away from the central pore. Some, however, belong to the six glutamate residues forming the selectivity filter, particularly their carboxylate groups. Visual inspection revealed no major alterations to the overall channel structure or pore geometry. The coordinate axes in Figure 2.1 illustrate the orientation of the transformed structure, with the pore axis aligned with the *z*-axis and the positive *z*-direction pointing toward the extracellular side.

## 3. Calcium channel triangular surface mesh

We generate triangular surface meshes for CalC6_3 using the GICMSG package [18], the repaired structure reported in 9DZW.pqr, and the following bounded hollow cylindrical domain,

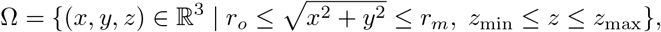

where *r*_*o*_ = 2, *r*_*m*_ = 48.393, *z*_min_ = −43.577, and *z*_max_ = 39.885, which we set to enclose the vdW surface of the CalC6_3 calcium channel. Here *r*_*o*_ is used to control the openness of the selectivity filter of an ion-selective channel protein. The hollow cylindrical domain Ω leaves the inner cylinder open while tightly enclosing the molecular structure of CalC6_3 toward the channel pore axis (the *z*-axis).

Because the channel pore is aligned with the *z*-axis, we transform Ω to a rectangular parameter domain, 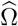, in the cylindrical coordinates (*r, θ, z*) as

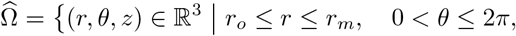

Note that 0 is excluded from the range of *θ* to avoid the duplication of points represented by *θ* = 0 and *θ* = 2*π*.

We partition the angular, axial, and radial intervals into 40, 80, and 60 subintervals, respectively. In GICMSG, the decay parameters *d*_*p*_ and *d*_*m*_ control the smoothness of the channel pore surface and the exterior molecular surface, respectively. A large value of *d*_*p*_ can produce a rugged channel pore surface, whereas reducing *d*_*m*_ can improve the smoothness of the exterior molecular surface. We performed two numerical tests using the default parameter values, except that *d*_*p*_ = 3.5 and *d*_*m*_ = 0.5 in Case 1 and *d*_*p*_ = 3.0 and *d*_*m*_ = 0.2 in Case 2. The resulting Gaussian triangular surface meshes provide two choices for the boundary ∂*D*_*p*_ of the protein region *D*_*p*_. Each mesh contains 9,998 vertices and 20,000 triangles, since we set the maximum allowed number of triangles, another parameter of GICMSG, to 20,000.

Figure 3.1 compares the two meshes in the top, side, bottom, and clipped views. The channel pore surfaces are highlighted in brown for clarity. In both cases, the top and bottom views show that the channel pore is open at both ends, while the clipped views reveal the pore surface along the *z*-axis. The comparison shows that Case 2 largely retains the channel pore geometry of Case 1 while producing a substantially smoother exterior molecular surface. These results demonstrate that GICMSG can achieve a good balance between preserving channel pore geometry and smoothing the exterior molecular surface.

**Fig. 3.1:**
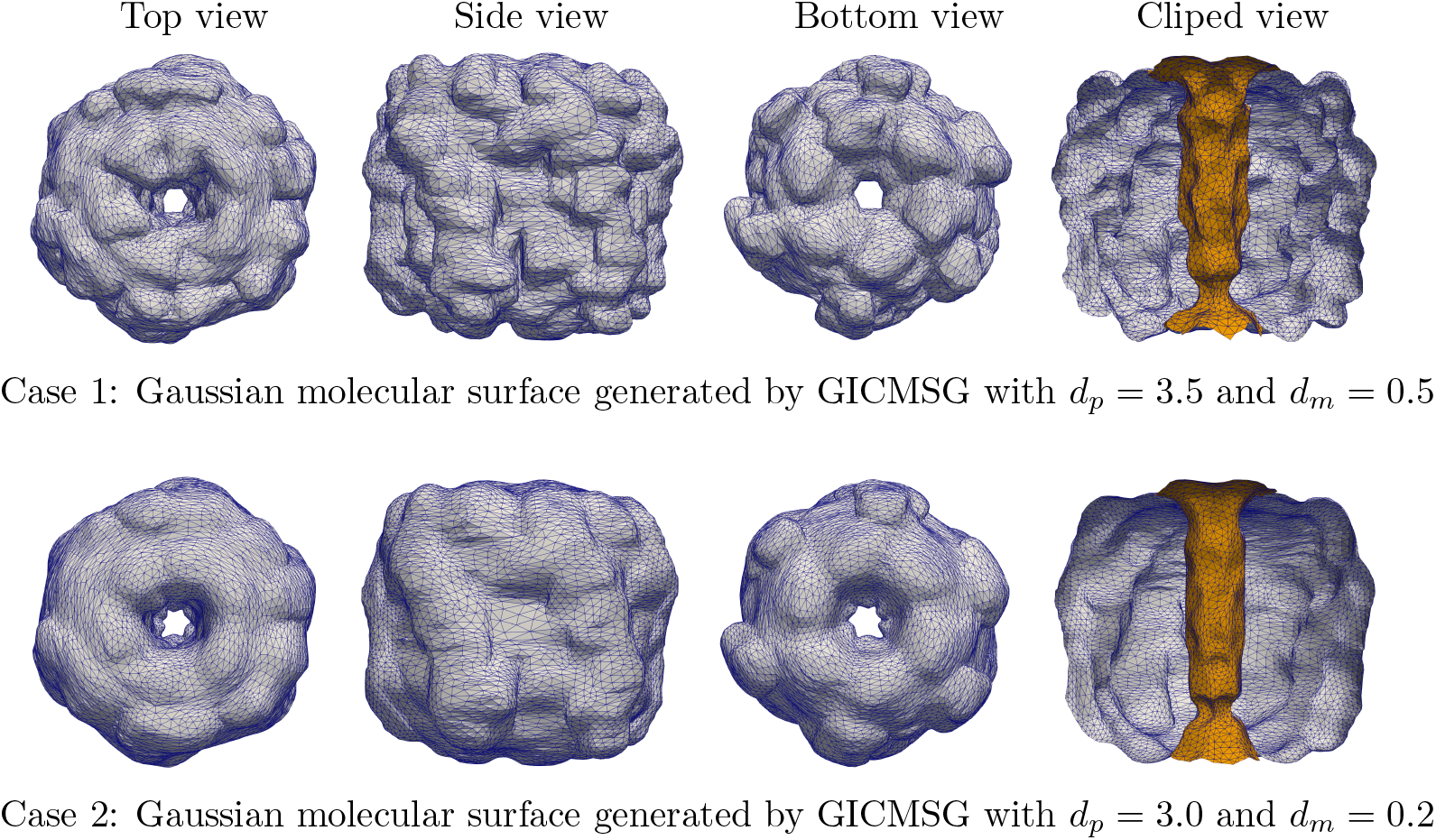
Comparison of the Gaussian triangular surface meshes generated by GICMSG in Cases 1 and 2. The channel pore surfaces are highlighted in brown.

To illustrate that the triangular surface mesh completely encloses the molecular structure of the CalC6_3 calcium channel, we use the mesh from Case 1, which wraps the structure more tightly than that from Case 2. Figure 3.2 compares this mesh with the molecular structure shown in sphere, line, and cartoon representations.

**Fig. 3.2:**
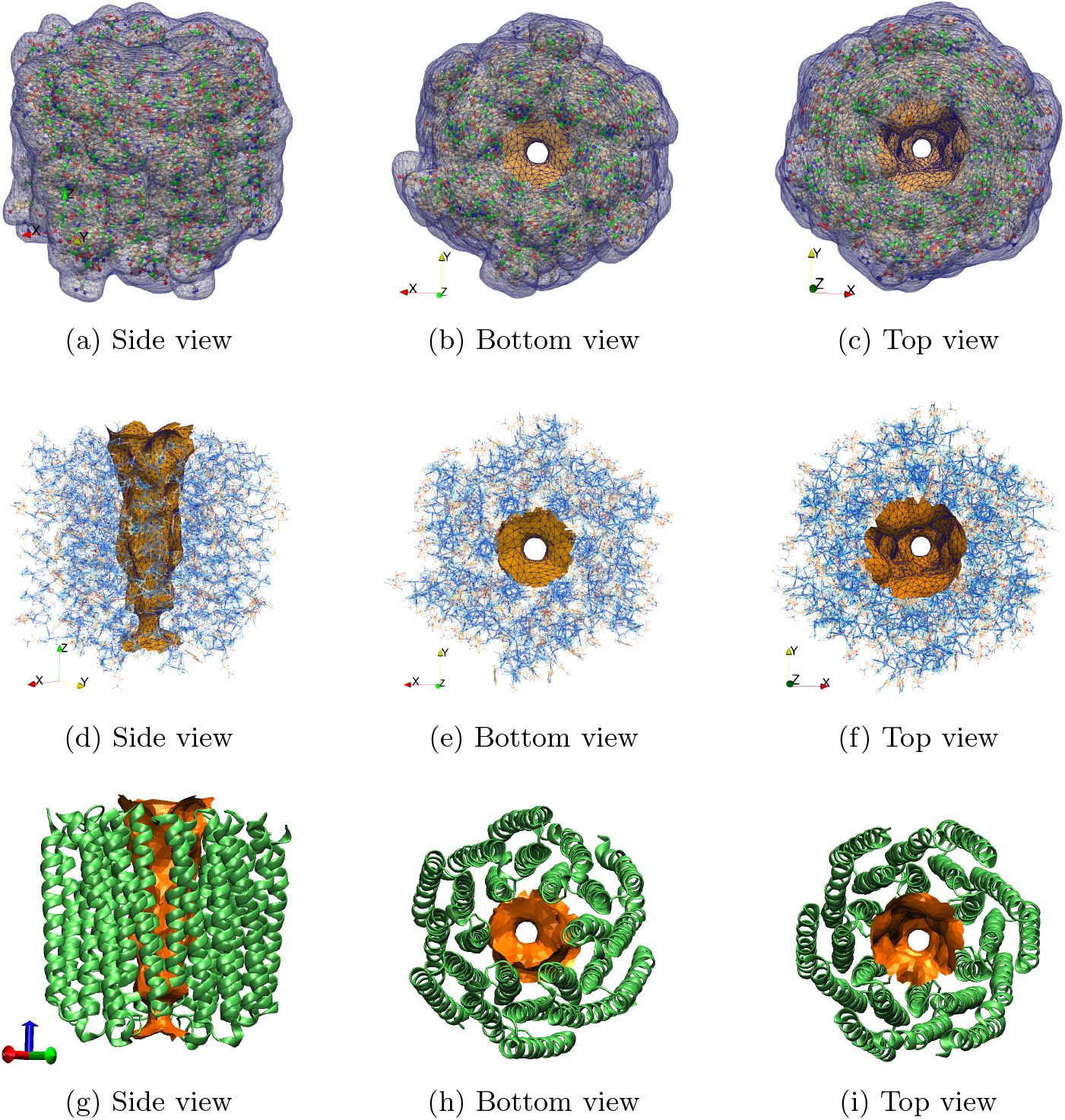
Comparison of the Gaussian triangular surface mesh ∂*D*_*p,h*_ of Case 1 with the CalC6_3 structure shown in (a)–(c) sphere, (d)–(f) line, and (g)–(i) NewCartoon representations. The channel pore surface is highlighted in brown.

Figure 3.2(a)–(c) shows that the triangular surface mesh of Case 1 encloses the molecular structure, with all 13,320 channel atoms represented as spheres. No atoms lie within the pore highlighted in brown or outside the exterior molecular surface.

Figure 3.2(d)–(f) compares the pore surface alone with the surrounding molecular structure shown in the line representation. The side view reveals the shape of the pore, while the bottom and top views show its geometry at the two openings. These views show that the channel atoms surround the pore surface without extending into the pore interior, leaving an atom-free channel pore across the cell membrane.

Figure 3.2(g)–(i) shows the relationship between the pore surface and the calcium channel structure in VMD’s NewCartoon representation. The side view illustrates the channel pore through the transmembrane helices, while the bottom and top views display the arrangement of the channel-forming helices around the central pore. Hence, Figure 3.2 demonstrates that the surface mesh encloses the molecular structure while preserving the channel pore geometry formed by the transmembrane helices.

## 4. Tetrahedral meshes of the simulation box

We selected the simulation box Ω = (−55, 55) × ( −56, 56) × (−62, 58) and placed the lower and upper membrane surfaces at *Z*1 = −16 and *Z*2 = 16, respectively. Here *Z*1 and *Z*2 are the two membrane location numbers used in our nuSMPBic model. Using the two Gaussian molecular surface meshes generated by GICMSG in Cases 1 and 2, we constructed two interface-fitted, unstructured tetrahedral meshes of Ω with the mesh-generation package reported in [9]. In each case, the box mesh Ω_*h*_ consists of the protein region mesh *D*_*p,h*_, the membrane region mesh *D*_*m,h*_, and the solvent region mesh *D*_*s,h*_. Here *D*_*p,h*_ is enclosed by the surface mesh ∂*D*_*p,h*_. The mesh data for both cases are reported in Table 4.1. Views of Ω_*h*_, *D*_*p,h*_, *D*_*m,h*_, and *D*_*s,h*_ for Case 1 are shown in Figure 4.1.

**Table 4.1:** Tetrahedral mesh data for Cases 1 and 2.

| Case | $D_{p,h}$ | $D_{m,h}$ | $D_{s,h}$ | $\Omega_h$ |
| --- | --- | --- | --- | --- |
| Number of mesh vertices |  |  |  |  |
| 1 | 41,601 | 14,354 | 22,980 | 65,874 |
| 2 | 44,535 | 13,264 | 24,193 | 69,336 |
| Number of tetrahedra |  |  |  |  |
| 1 | 233,861 | 61,726 | 104,327 | 399,914 |
| 2 | 252,478 | 57,270 | 110,919 | 420,667 |

**Fig. 4.1:**
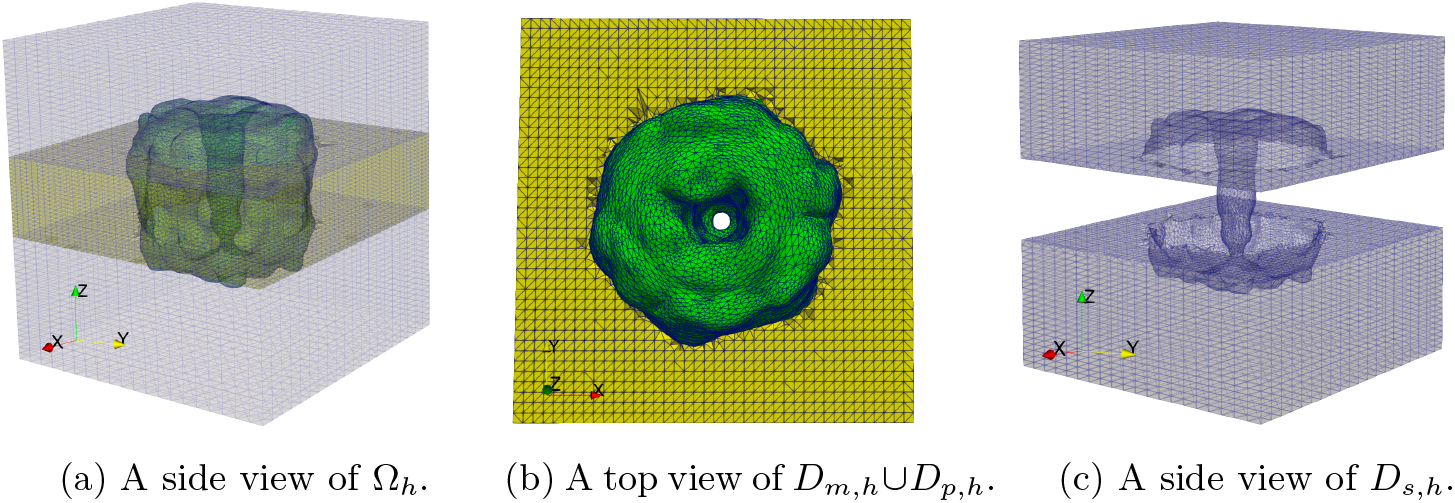
The box domain mesh Ω_*h*_, the protein region mesh *D*_*p,h*_, the membrane region mesh *D*_*m,h*_, and the solvent region mesh *D*_*s,h*_. In (a, b), *D*_*p,h*_ and *D*_*m,h*_ are highlighted in green and yellow, respectively. In (a), *D*_*s,h*_, *D*_*p,h*_, and *D*_*m,h*_ are shown in low opacity to expose *D*_*p,h*_, *D*_*m,h*_, and the channel pore.

Figure 4.1(a), (b), and (c) presents a side view of Ω_*h*_, a top view of *D*_*m,h*_ and *D*_*p,h*_, and a side view of *D*_*s,h*_, respectively. These views illustrate the geometry and openness of the channel pore, as well as the complex interfaces between the protein, membrane, and solvent regions. The pore is funnel-shaped, with a wide extracellular entrance that gradually narrows to a slender, approximately cylindrical selectivity filter before opening into a wider solvent-filled cavity toward the intracellular side.

## 5. Numerical test results and discussion

To investigate the calcium selectivity of the CalC6_3 channel, numerical experiments were performed using our integrated package. The ionic solution consists of 0.1 M NaCl and 0.1 M CaCl_2_. Accordingly, the bulk concentrations of Ca^2+^, Na^+^, and Cl^−^ are 0.1, 0.1, and 0.3 mol/L, respectively, satisfying electroneutrality. We explored the effects of ion hydration by using hydrated radii of Ca^2+^, Na^+^, and Cl^−^ ions as 4.12, 3.58, and 3.32 Å, respectively, which come from the BioNumbers database (https://bionumbers.hms.harvard.edu/bionumber.aspx?id=108517). All other numerical parameters were set to their default values in the nuSMPBic package.

The numerical experiments were conducted on a single CPU core of a Mac Studio equipped with an Apple M4 Max processor and 128 GB of memory. Table 5.1 reports the CPU times and the numbers of iterations required to satisfy the iteration termination criterion ∥*F* ∥_2_ *<* 0.001, where ∥*F*∥_2_ denotes the Euclidean norm of the residual vector of the nonlinear finite element system for the nuSMPBic model. The numerical results are shown in Figures 5.1 and 5.2.

**Table 5.1:**
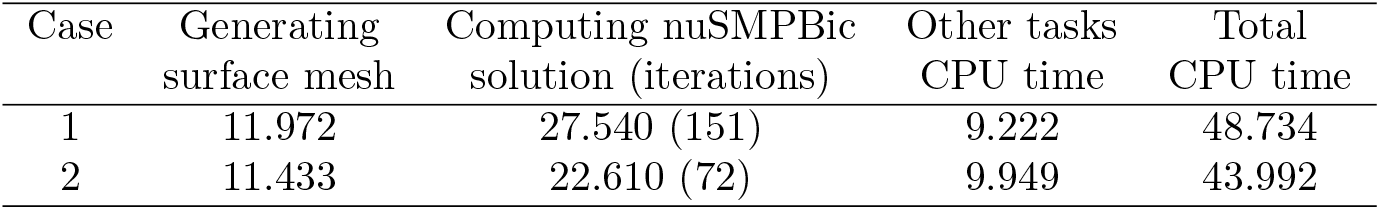
CPU times (in seconds) for Cases 1 and 2. Other tasks include input data reading, tetrahedral mesh generation, and output data saving. Numbers in parentheses are the numbers of iterations.

| Case | Generating<br>surface mesh | Computing nuSMPBic<br>solution (iterations) | Other tasks<br>CPU time | Total<br>CPU time |
| --- | --- | --- | --- | --- |
| 1 | 11.972 | 27.540 (151) | 9.222 | 48.734 |
| 2 | 11.433 | 22.610 (72) | 9.949 | 43.992 |

**Fig. 5.1:**
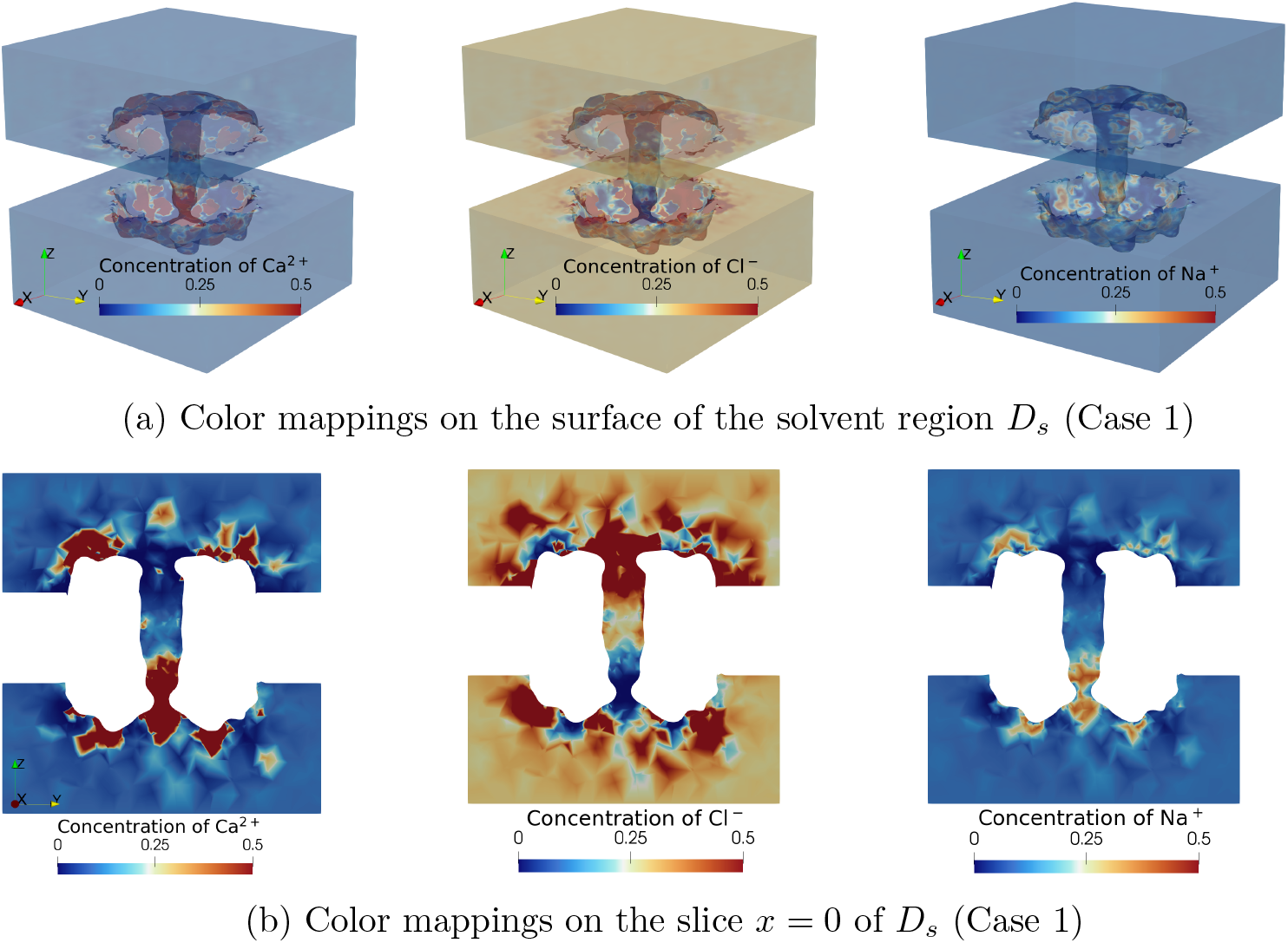
A comparison of the concentration of Ca^2+^ with the concentrations of Cl^−^ and Na^+^ in color mapping for the CalC6_3 calcium channel (PDB ID: 9DZW) in an ionic solution of 0.1 molar CaCl_*s*_ and 0.1 molar NaCl: (a) mappings on the surfaces of the solvent region *D*_*s*_ and (b) mappings on the cross-section *x* = 0 of *D*_*s*_.

**Fig. 5.2:**
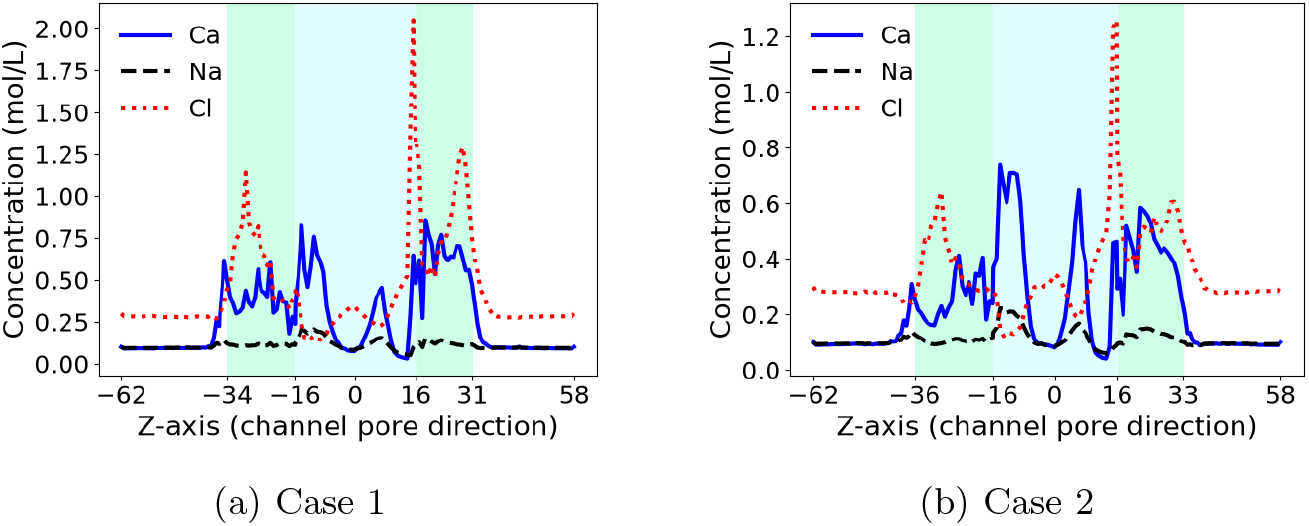
Comparisons of the concentration functions of Ca2+, Cl^−^, and Na^+^ of Case 1 with those of Case 2 in block-averaged curves along the channel pore (*z*-axis) direction. Here, the pore part merged in the membrane ( −16 ≤ *z ≤* 16) is shaded light-cyan, and the other parts of the pore are shaded green.

### 5.1. Performance of the integrated software package

As reported in Table 4.1, Ω_*h*_ and *D*_*s,h*_ contain 65,874 and 22,980 vertices in Case 1 and 69,336 and 24,193 vertices in Case 2, respectively. In the nuSMPBic finite element solver, the finite element approximations of the electrostatic potential function *u* and the three ionic concentration functions are defined on Ω_*h*_ and *D*_*s,h*_, respectively, and computed by solving a nonlinear finite element system. Thus, the nonlinear systems for Cases 1 and 2 contain 134,814 and 141,915 unknowns, respectively, before accounting for boundary conditions.

Table 5.1 demonstrates the efficiency of the nuSMPBic finite element solver and the high performance of the integrated software package. The numerical experiments for Cases 1 and 2 were completed in about 49 and 44 seconds, respectively, on a single CPU core, including about 12 and 11 seconds for Gaussian molecular surface mesh generation using GICMSG. Solving the nonlinear system in Case 2 required 72 iterations, compared with 151 in Case 1, while the computing time decreased from about 28 to 23 seconds, despite the larger nonlinear system in Case 2. These results suggest that a smoother molecular surface mesh can improve the convergence and performance of the nuSMPBic finite element solver.

### 5.2. Color mappings of ionic concentration functions

Because the distribution patterns of ions in Case 2 are similar to those in Case 1, we only consider the ionic concentration functions derived from Case 1 in Figure 5.1.

Figure 5.1 compares the concentrations of Ca^2+^, Cl^−^, and Na^+^ in color mapping onto the surfaces of the solvent region *D*_*s*_ in Figure 5.1(a) and onto the cross-section *x* = 0 of *D*_*s*_ in Figure 5.1(b). They all use the same color scale with blue for 0 and red for the concentration values greater than 0.5 mol/L. From these color mappings, it can be seen that the three ionic species exhibit markedly different concentration patterns. Ca^2+^ accumulates strongly near the intracellular entrance and in the adjacent portion of the channel pore. In contrast, Na^+^ is largely excluded from the pore, although it has the same bulk concentration of 0.1 mol/L as Ca^2+^ and a smaller hydrated radius than that of Ca^2+^. Thus, ion size alone cannot explain the competition between these two cations for the limited space inside the pore. Instead, the charge number plays a major role in fighting for space within the narrow channel pore. Competition among charged ions for space plays a major role in the crowded channel pore. Furthermore, the preferential accumulation of Ca^2+^ over Na^+^ near the intracellular entrance provides (b) Color mappings on the slice *x* = 0 of *D*_*s*_ (Case 1) evidence that this region contributes to calcium recognition. The role of crowding near the intracellular entrance has received limited attention in the current literature.

Figure 5.1 also shows that the concentration of Cl^−^ exhibits a different distribution pattern from the concentrations of Ca^2+^ and Na^+^. In particular, Cl^−^ ions accumulate mainly near the extracellular entrance and in portions of the surrounding solvent region, while its concentration is low in parts of the pore where Ca^2+^ accumulates. It should also be noted that the bulk concentration of Cl^−^ is 0.3 mol/L, which is three times the bulk concentration of either cation. The distinct distributions of Ca^2+^, Na^+^, and Cl^−^ reflect the combined effects of ionic charge, ionic size, and the heterogeneous electrostatic environment created by the channel.

### 5.3. Block-averaged profiles of ionic concentration functions

Color mappings of ionic concentration functions show only the function values on a surface or cross-section of the solvent region *D*_*s*_, and concentration values above 0.5 mol/L cannot be distinguished because of saturation of the color scale. To provide a quantitative description of ionic concentrations throughout *D*_*s*_, we generated block-averaged profiles using the two-dimensional (2D) curve mapping scheme reported in [17]. Specifically, we divided the *z*-axis interval [−62, 58] into subintervals, [−62, −34], [−34, −16], [−16, 16], [16, 31], and [31, 58], and divided each subinterval uniformly into 20 parts. This partitions *D*_*s*_ into 100 blocks. We then calculated the average concentrations of Ca^2+^, Na^+^, and Cl^−^ over each block, obtaining 100 points for each concentration function. Using these points, we plotted three 2D curves as functions of *z* for both Case 1 and Case 2. Note that the *z*-axis coincides with the channel pore axis and the membrane-normal direction. Hence, these curves characterize the block-averaged ionic concentration profiles across the membrane and throughout the solvent region. The resulting curves are displayed in Figure 5.2. For clarity, the portion of the pore within the membrane, −16 ≤ *z* ≤ 16, is highlighted in light cyan. The two adjacent pore portions, −34 ≤ *z* ≤ −16 and 16 ≤ *z* ≤ 31, are highlighted in green.

### 5.4. Ionic concentration profiles along the channel pore

Figure 5.2 compares the block-averaged ionic concentration profiles of Cases 1 and 2. Both cases exhibit similar profiles along the channel pore, even though the exterior Gaussian molecular surface mesh in Case 2 is much smoother than that in Case 1. This agreement suggests that smoothing the exterior molecular surface preserves the main ionic concentration patterns in these tests. In both cases, the Ca^2+^ concentration substantially exceeds its bulk value of 0.1 mol/L in several parts of the pore, reaching approximately 0.8 mol/L near *z* = −14. It remains elevated over much of the intracellular pore region. Moreover, Ca^2+^ is strongly enriched, while the Cl^−^ concentrations are substantially reduced relative to their bulk values within −16 ≤ *z ≤ −*7, where the calcium selectivity filter formed by six Glu residues is located. The predominantly negative electrostatic environment in this region favors cation accumulation and suppresses anion accumulation.

In both cases, the Na^+^ concentration remains close to its bulk value of 0.1 mol/L over most of the channel pore and exhibits only modest local increases. The Cl^−^ concentration, however, exhibits a distinctly different profile because its bulk value of mol/L is three times that of either cation. Its largest block-averaged concentration occurs near *z* = 16, at the upper membrane surface. Two other peaks appear near the intracellular and extracellular entrances of the channel pore, at approximately *z* = −28 and *z* = 30, respectively. It is important to note that the Cl^−^ bulk concentration is 0.3 mol/L, three times that of either cation.

The preferential accumulation of Ca^2+^ over Na^+^, despite their equal bulk concentrations and the larger hydrated radius assigned to Ca^2+^, indicates that hydrated ion size alone does not determine ionic accumulation within the pore. The results highlight the importance of ionic valence and the channel’s molecular geometry and charge distribution. This preferential accumulation is consistent with the experimentally reported calcium selectivity of CalC6_3 [12], showing that the nuSMPBic model captures an equilibrium feature relevant to calcium selectivity in this channel.

## 6. Conclusions

This work has developed an integrated software package for computational studies of the de novo designed calcium channel CalC6_3. The package combines molecular structure preparation, Gaussian Ion Channel Molecular Surface Generation (GICMSG), interface-fitted mesh generation, and the nonuniform size-modified Poisson–Boltzmann ion channel (nuSMPBic) finite element solver. This integration streamlines the complete workflow from structural data preparation to mesh generation and numerical solution.

Using the integrated package, the original 9DZW PDB file is cleaned, repaired, and aligned with the coordinate system required by the nuSMPBic model. A corresponding PQR file provides the atomic coordinates, partial charges, and radii needed for the simulations. GICMSG then generates a triangular molecular surface mesh that preserves the intricate channel pore geometry while maintaining a sufficiently smooth exterior molecular surface. This surface mesh is used to construct a high-quality, interface-fitted tetrahedral mesh of the simulation box consisting of the protein, membrane, and solvent regions.

Numerical experiments have been performed using two Gaussian molecular surfaces generated by GICMSG, one of which has a much smoother exterior surface than the other. The test results confirm preferential Ca^2+^ accumulation in the narrow selectivity filter formed by six Glu residues, supporting the experimentally observed calcium selectivity of CalC6_3. The complete numerical calculation, including solving a nuSMPBic finite element system with 141,915 unknowns, required only about 44 seconds on a single CPU core of a Mac Studio equipped with an Apple M4 Max processor, demonstrating the computational efficiency of the integrated package. The package thus provides a useful computational tool for studying CalC6_3 and can also be adapted to simulate other ion channel proteins. A detailed description of its construction and usage as a general tool for ion channel protein simulations, particularly of selectivity, will be presented in a separate paper.

## Acknowledgments

This work was partially supported by the National Science Foundation, USA, through award number DMS-2153376, and the Simons Foundation, USA, through record ID MPS-TSM-00711776.

## Notes

### Competing Interest Statement

The authors have declared no competing interest.

